# Genetic regulators of neuronal survival across metabolic environments

**DOI:** 10.64898/2025.12.19.695350

**Authors:** Neal Bennett, Yanilka Soto-Muniz, Jonathan X. Meng, Megan Lee, Joyce Yang, Will R Flanigan, Alexander R. Pico, Isha H Jain, Ken Nakamura

## Abstract

Cellular energy metabolism and oxygen availability shape neuronal function and vulnerability, yet the genetic regulators of these metabolic processes in human neurons remain incompletely understood. Here, we performed CRISPR interference (CRISPRi) screens in human induced pluripotent stem cell (iPSC)-derived neurons across four distinct metabolic conditions and at three physiologically relevant oxygen tensions. This combinatorial approach enabled systematic interrogation of gene-environment interactions that govern neuronal metabolic adaptation. We identified genes—including genes associated with Leigh syndrome and autism spectrum disorder—whose importance for cell survival is highly sensitive to environmental context, revealing potential mechanisms underlying metabolic specification and selective neuronal vulnerability in neurological disorders. Our screens also uncovered regulators of neuronal glycolysis, including *KIAA1429* and *MAPT* among others, which are previously uncharacterized modulators of neuronal glucose utilization and metabolic flexibility. Our work nominates candidate metabolic interventions and gene targets for enhancing neuronal resilience under hypoxic or nutrient-limited conditions.

## INTRODUCTION

ATP fuels essentially all cellular functions. The primary cellular processes that generate ATP in mammalian cells—glycolysis, the citric acid cycle and oxidative phosphorylation—are well characterized. However, energy production depends on both the availability of energy fuels and substrates such as glucose and oxygen, and on the integrity of the different steps of energy production, which can vary widely in physiologic and disease states. For example, brain glucose uptake rises sharply in early childhood and then declines with age [1, 2]. In cancer, and potentially Alzheimer’s disease and Parkinson’s disease as well, cells shift towards aerobic glycolysis, where a larger proportion of glucose is metabolized through glycolysis or other non-oxidative metabolic pathways than through oxidative phosphorylation [3–5]. The cellular basis for these metabolic shifts is poorly understood.

Considerable evidence indicates that neurons produce energy under physiological conditions from two main sources, lactate supplied by astrocytes and glucose taken up directly and metabolized through glycolysis, with both substrates converted to pyruvate and oxidized in the citric acid cycle [6, 7]. However, we lack a comprehensive understanding of how neurons regulate these critical energy pathways, and how genetic deficiencies and environmental nutrients can combine to disrupt neuronal development or promote degeneration. As such, we also don’t know whether mismatches between neurons’ energy needs and energy supplies might be improved by targeted metabolic interventions, or whether such interventions would be neuroprotective.

To begin to address these gaps, we performed parallel gene-by-environment CRISPR-based screens aimed at delineating the influence of energy fuels and substrates on neuronal survival, and at systematically identifying regulators of essential metabolism genome-wide.

## RESULTS

### Gene-by-environment screens for regulators of neuronal survival

We expressed genome-wide CRISPRi sgRNA libraries [8] in human neurons generated from induced pluripotent stem cells (iPSC) [9, 10], and quantified the impact of gene knockdown on neuronal survival as a function of energy substrates. Specifically, 18-day-old neurons were incubated in three types of media: (1) allowing both respiration and glycolysis, (2) forcing the cells to rely on glycolysis, and (3) forcing them to rely on respiration for energy. These three conditions model metabolic extremes that may occur in disease [11, 12], and we’ve shown previously that they lead to reliance on distinct genes for energy production [13]. A fourth metabolic condition was designed to mimic physiologic levels of brain energy substrate and oxygen (1.5 mM glucose, 200 µM pyruvate, 0.2 mM beta-hydroxybutyrate, 1 mM lactate and750 µM glutamine, in 4% O_2_ [14–21]).

Oxygen is essential for hundreds of reactions including aerobic ATP production [22, 23]. Since oxygen levels in the brain (1-5%[19]) are markedly lower than in room air (20%), and can further decrease in ischemia, we also assessed all four energy substrate conditions at three distinct oxygen levels (20%, 5%, 0.3%), for a total of 12 experimental conditions.

After three days in each condition, neurons were harvested, and CRISPRi sgRNAs were amplified and sequenced. Survival phenotypes were calculated based on log-fold changes in sgRNA abundance after versus before treatment (Fig 1A).

**Fig 1.**
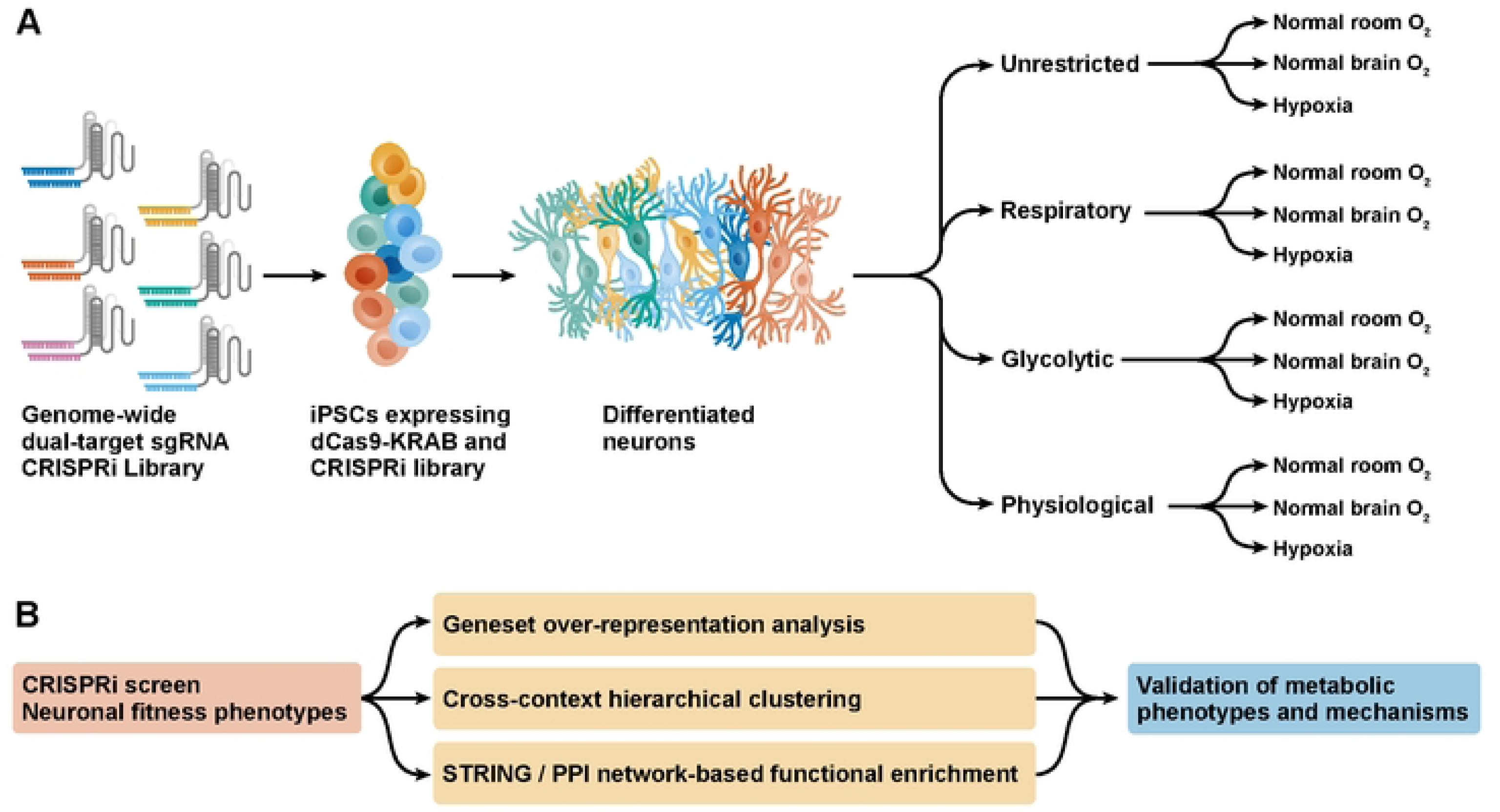
A CRISPRi screen in human induced pluripotent stem cell (iPSC)-derived neuronal cells across metabolic states. A) We expressed a genome-wide dual-sgRNA CRISPRi library in human iPSCs. iPSCs were differentiated into neuronal cells for 18 days before coercion for 3 days into one of 12 metabolic states corresponding to 4 culture media and 3 oxygen tensions. The culture media 1) provided ample substrates for both glycolysis and oxidative phosphorylation (“unrestricted”), 2) limited energy metabolism respiration only (“Respiratory”) or 3) to glycolysis only (“Glycolytic”), or 4) approximated the metabolite concentrations observed in the physiologic brain (“Physiological”). The oxygen fraction in the culture chambers was either 20% (“Normal Room O_2_”), 5% (“Normal Brain O_2_”), or 0.3% (“Hypoxia”). The screen was conducted in n = 2 replicates. B) To analyze data from the screen, we employed multiple bioinformatics strategies examining functional enrichment. We looked for patterns in biological and disease-related gene groups across individual specific metabolic states and across broad groupings of similar metabolic states, mapped known interactions between survival-related genes, and used clustering to find new groups of genes with similar functions. Finally, we validated and tested mechanisms for metabolic phenotypes in individual gene knockdown lines.

### Mitochondrial genes are essential for respiration, while the pentose phosphate pathway is critical for glycolysis

To analyze the results of these parallel screens, we combined geneset over-representation analysis, network-based functional enrichment, and unbiased hierarchical clustering (Fig 1B).

Through geneset over-representation analysis, we identified distinct genetic pathways that regulated neuronal survival when cells were forced to rely on glycolysis only or respiration only across oxygen levels. As expected, knockdown of genes associated with the biological process of cellular respiration, and most prominently genes with roles in mitochondrial complex I assembly, was highly lethal when analyzed across aggregated respiration-only metabolic states (NDUFA8, NDUFB9, NDUFV1, among others, adjusted p = 0.028, Fig 2A, Table S1). When considering individual respiratory-only conditions, mitochondrial pathways were generally more essential at higher oxygen levels. However, genes associated with cellular respiration and mitochondrial complex I assembly were depleted in both the high- and low-oxygen respiratory conditions, presumably because mitochondrial respiration is required for neuronal survival in that setting (Aerobic electron transport chain: NDUFB4, UQCRC2, CYC1, among others, adjusted p = 0.001058 in 20% oxygen, adjusted p = 0.006521 in 0.3% oxygen, Complex I biogenesis: NDUFA8, NDUFB9,NDUFV1, among others, adjusted p = 0.0000026 in 20% oxygen, p = 0.000435 in 0.3% oxygen, Fig 2B, Table S2)

**Fig 2.**
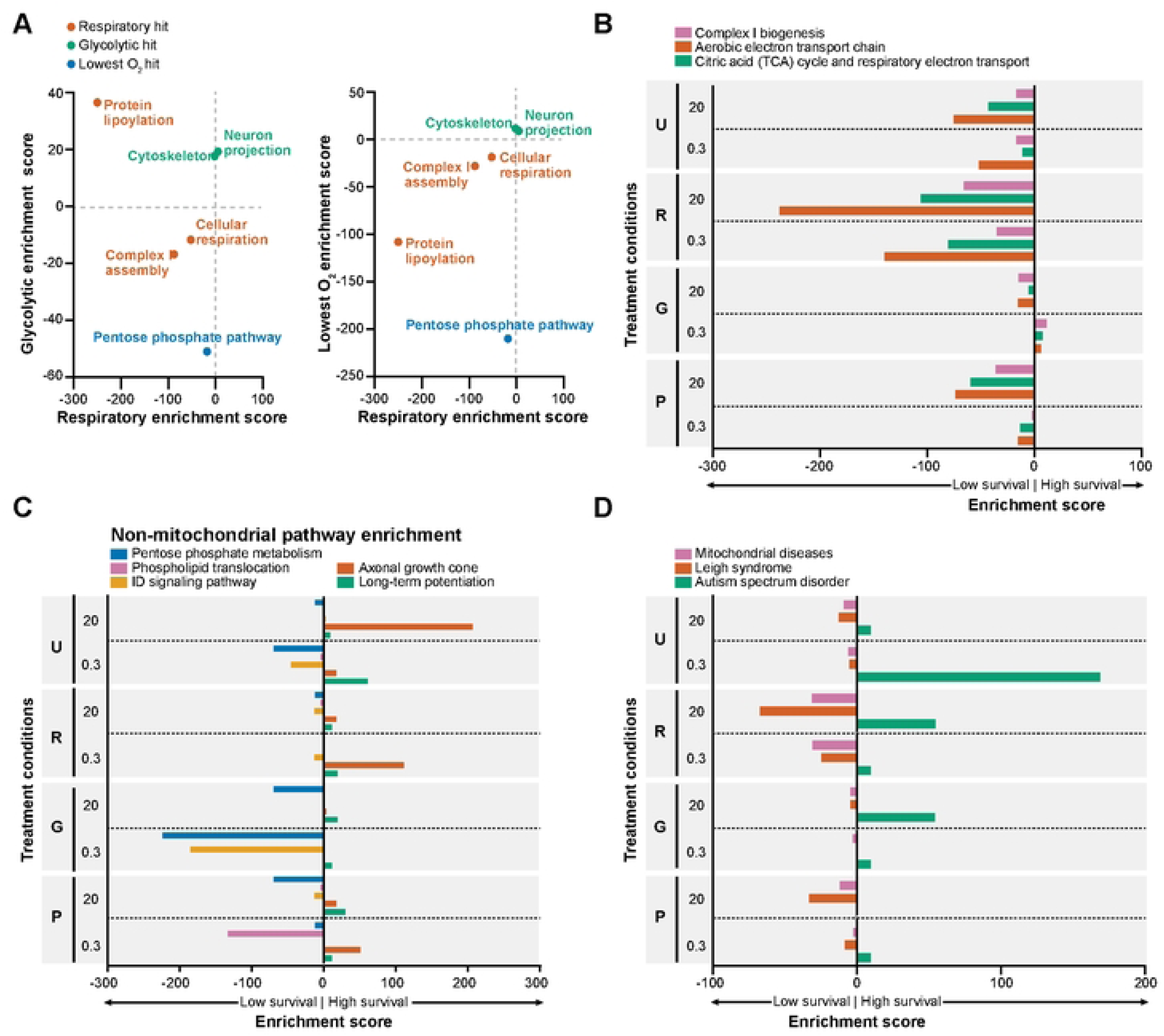
Identification of genetic pathways that regulate neuronal survival in individual and aggregated metabolic states. A) Over-representation analysis among the gene KDs with the lowest average survival across averaged Respiratory states reveals the essentiality of genes involved in cellular respiration, complex I assembly, and protein lipoylation. Among gene KDs that limit survival across averaged Glycolytic states, we observe an over-representation of genes that code for parts of the cytoskeleton and neuron projection proteins. Genes encoding the pentose phosphate pathway are essential across averaged Hypoxia metabolic states. All “hit” pathways shown had an adjusted p < 0.05 in their respective aggregated metabolic state. B–D) Over-representation analysis among the genes with the greatest impact on survival in each individual metabolic state reveals genetic pathways that are either mitochondrial (B), non-mitochondrial (C), or (D) disease- and disorder-associated genes. Metabolic states were set by using media that enabled either 1) unrestricted respiration and glycolysis: “U”, 2) respiration only: “R”, 3) glycolysis only: “G”, or 4) contained physiological levels of respiratory and glycolytic fuels: “P”. All displayed pathways had an adjusted p < 0.05 in at least one individual metabolic state.

In contrast, the glycolytic pathway was not essential under any condition, even when neurons were forced to rely on glycolysis for energy. This finding may seem surprising, but it aligns with our previous observations that knocking down individual glycolytic genes often did not impact glycolytic ATP[13], presumably because of the redundancy of many glycolytic enzymes. Interestingly, knockdown of genes in the pentose phosphate pathway was associated with decreased survival in both glycolysis-only and low-oxygen conditions (PRPS1, G6PD, PGD, among others, Fig 2A). This could reflect a requirement for redox management or biosynthetic products of the PPP under these metabolic states. Consistent with this hypothesis, cells with knockdown of genes associated with the pentose phosphate pathway were most depleted specifically in conditions of forced glycolysis and low oxygen exposure (RPE, PGLS, PGD, Fig 2C).

Interestingly, cells with knockdown of genes that encode cytoskeleton proteins (NF2, EZR, TUBB1 among others) or neuron projection proteins (SHANK2, SYT1, GPM6A among others) were positively selected in glycolysis-only conditions (Fig 2A, Table S1). While genes that code for neuron projection were not significantly enriched in any individual metabolic state, cells with knockdown of genes encoding for the axonal growth cone (L1CAM, DCC, FLRT3 among others), a specialized neuronal structure, were enriched in a range of metabolic states, particularly when metabolism was unrestricted and oxygen was high. It has long been known that injured axons can “dedifferentiate” into axonal growth cones [24]. Further studies will need to determine whether loss of these genes contributes to altered neuronal metabolism by slowing the trajectory of differentiation or promoting metabolic plasticity as opposed to commitment to reliance on mitochondrial respiration.

### Loss of Leigh syndrome-associated genes leads to respiratory vulnerability while loss of autism spectrum disorder genes imparts metabolic flexibility

We next examined if the gene knockdowns we identified were associated with inherited metabolic diseases. Cells with knockdown of mitochondrial disease genes, and specifically genes whose mutations cause Leigh syndrome, were selectively vulnerable when forced to rely on respiration only in high oxygen (Leigh syndrome with leukodystrophy: NDUFV1, NDUFS6, UQCRC2 among others, enrichment in 20% oxygen respiratory conditions adjusted p-value = 0.0078, 5% oxygen adjusted p-value = 0.21, 0.3% oxygen adjusted p-value = 0.57, Fig 2D, Table S3). These observations are consistent with the fact that many of the genes associated with Leigh syndrome and mitochondrial diseases encode for components of the respiratory chain in mitochondria[25], that individuals suffering from these diseases demonstrate exercise intolerance, and that cells derived from Leigh syndrome patients have impaired aerobic respiration[13].

We were surprised to observe that cells with knockdown of certain genes associated with autism spectrum disorder (ASD), particularly PTCHD1 and SHANK2, led to enrichment in several metabolic states. Similar to what we observed with knockdown of genes involved in axonal growth cone function or long-term potentiation (Fig 2C), further studies will be needed to determine if ASD-associated mutations cause a loss-of-function that disrupts the metabolic specification normally accompanying neuronal differentiation.

### Protein association networks of neuron metabolic regulators nominate NEDDylation and β-arrestin signaling as hypoxia-sensitive processes

Some genetic pathways may contain multiple genes that strongly regulate neuron survival, but through different mechanisms. For that reason, we looked for protein partners of the strongest hits from any condition in the screen using STRING protein-protein interaction (PPI) annotation. This approach takes into consideration the independently accumulated data on the myriad ways our top genes interact and coordinate to perform biological functions.

Indeed, several of these survival-regulating genes had well-characterized associations with each other (Fig 3A, Table S4). For instance, genes involved in mitochondrial respiratory chain complex I assembly form a tight web of interactions and were generally essential for neuronal survival (cluster 6: NDUFAF6, NDUFV1, NDUFS6 among others, Fig 3A). Loss of gene expression from this cluster conferred a vulnerability in respiration-only conditions versus glycolytic conditions that was blunted in low-oxygen conditions (5/21 nodes with survival phenotypes < -5 log-fold change in respiration-only versus glycolysis-only in high oxygen, versus only 1/21 nodes with survival phenotypes < -5 log-fold change in low oxygen, Fig 3B).

**Fig 3.**
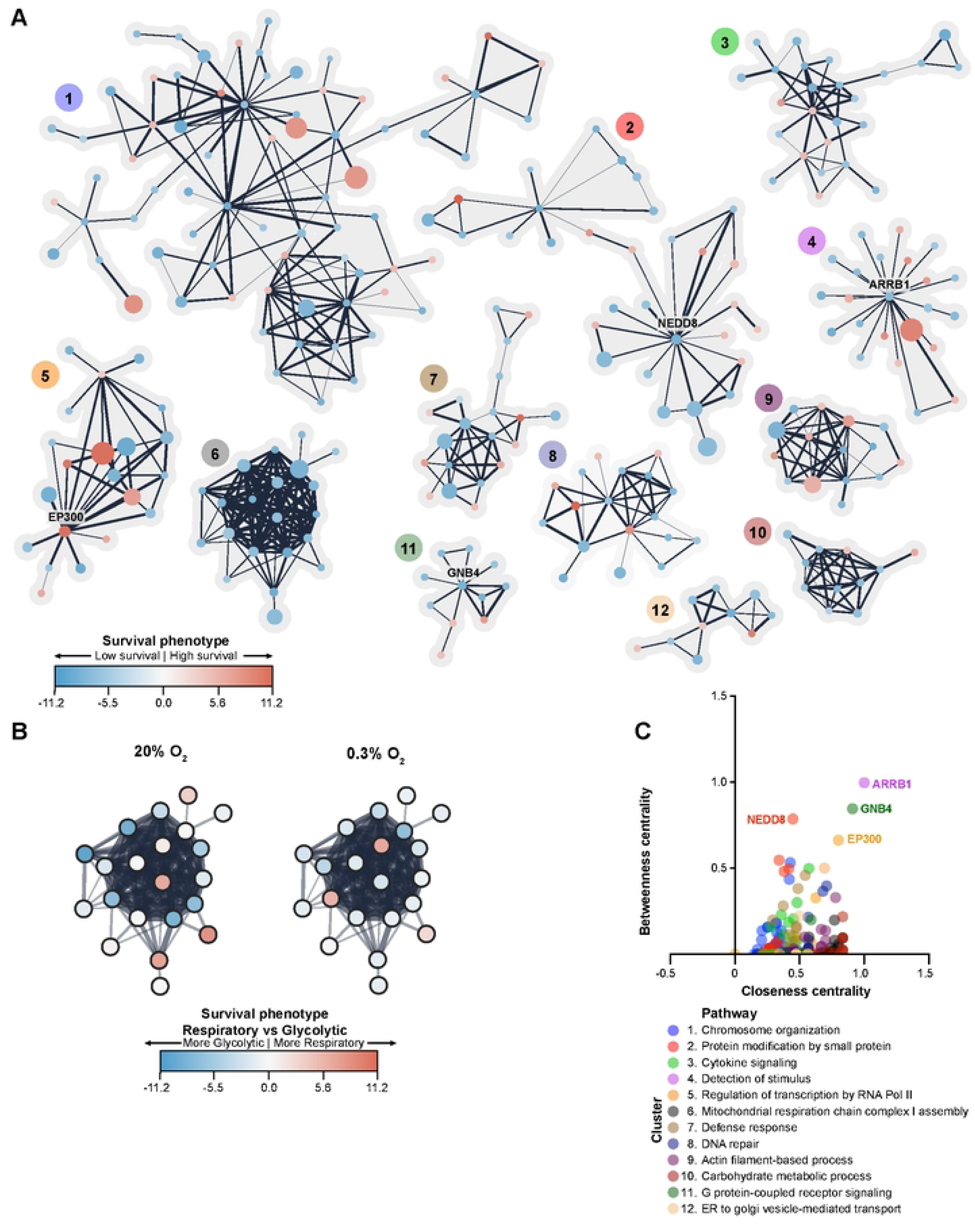
Network and STRING-based functional enrichment among neuronal metabolic survival genes. (A) Genes that regulated neuronal metabolic survival in at least one metabolic state were clustered using the Markov Cluster Algorithm (MCL). Each node in this network corresponds to one gene knockdown. Node color represents the maximal survival phenotype observed for this node’s KD across all metabolic states. Node size corresponds to the number of metabolic states in which the node’s KD significantly affected metabolic survival. Functional enrichment in network clusters was based on the STRING interaction database. (B) KD’ing genes in cluster 6 (enriched for mitochondrial respiratory chain complex I assembly genes) renders cells more vulnerable to respiration-only than glycolysis-only conditions at high oxygen (blue fill color indicates low survival). This difference is blunted in low oxygen (less intense blue node fills). (C) Network topological analysis reveals central nodes of neuronal metabolic regulation. Betweenness centrality measures how often a node lies on the shortest path between two other nodes. Closeness centrality measures how close a given node is to all other nodes.

The PPI network analysis allowed us to define genes that are central to the PPI clusters (Fig 3C). Central genes are defined as those most likely to be on the shortest path between two nodes of a network (“betweenness centrality”) and that display the shortest average distance to all other nodes in the network (“closeness centrality”). We found that central genes were associated with neuronal development or neuronal differentiation, mediated through direct conjugation (NEDD8)[26], regulation of transcription factors (EP300)[27], or by integrating signals between receptors and signaling effectors (GNB4, ARRB1)[28, 29]. Notably, NEDD8 and CDC34 are linked based on STRING data, likely due to their shared role in regulating the multi-protein E3 ubiquitin ligase Skp, Cullin, F-box containing complex[30]. Knockdown of these two genes resulted in similar effects on neuronal survival in multiple hypoxic conditions (Fig S1A). Further work will be needed to determine what action or ubiquitinated targets of this E3 ubiquitin ligase complex are critical for neuronal survival in hypoxia. ARRB1 (arrestin β-1) is known to limit olfactory receptor internalization and associated signaling[31]. There is growing appreciation for olfactory dysfunction as an early symptom of many neurodegenerative diseases[32, 33]. However, to our knowledge, a link between defects in specific olfactory receptors and associated signaling and neuronal metabolism has not been established. These results nominate specific interactions between β-arrestin and signaling from specific olfactory receptors (OR6K3, OR5M10) that could be suppressed to promote neuronal survival in hypoxic conditions (Fig S1A).

### Hierarchical clustering reveals broad essentiality of aerobic respiration genes in neuron survival

While complex I assembly genes were generally essential for neuronal survival, many glycolytic genes were not as consistently essential for neuronal survival (1/11 nodes with survival phenotypes < -2 in high oxygen, glycolysis-only conditions Fig S1B). While protein associations provide a powerful way to identify genes that may work in the same pathways, they may not reflect the neuron-specific cellular context, since PPI studies are not necessarily conducted in neurons. To identify functional similarity between essential genes in an alternative, unbiased way, we used a hierarchical clustering approach (Fig 4, Table S5).

**Fig 4.**
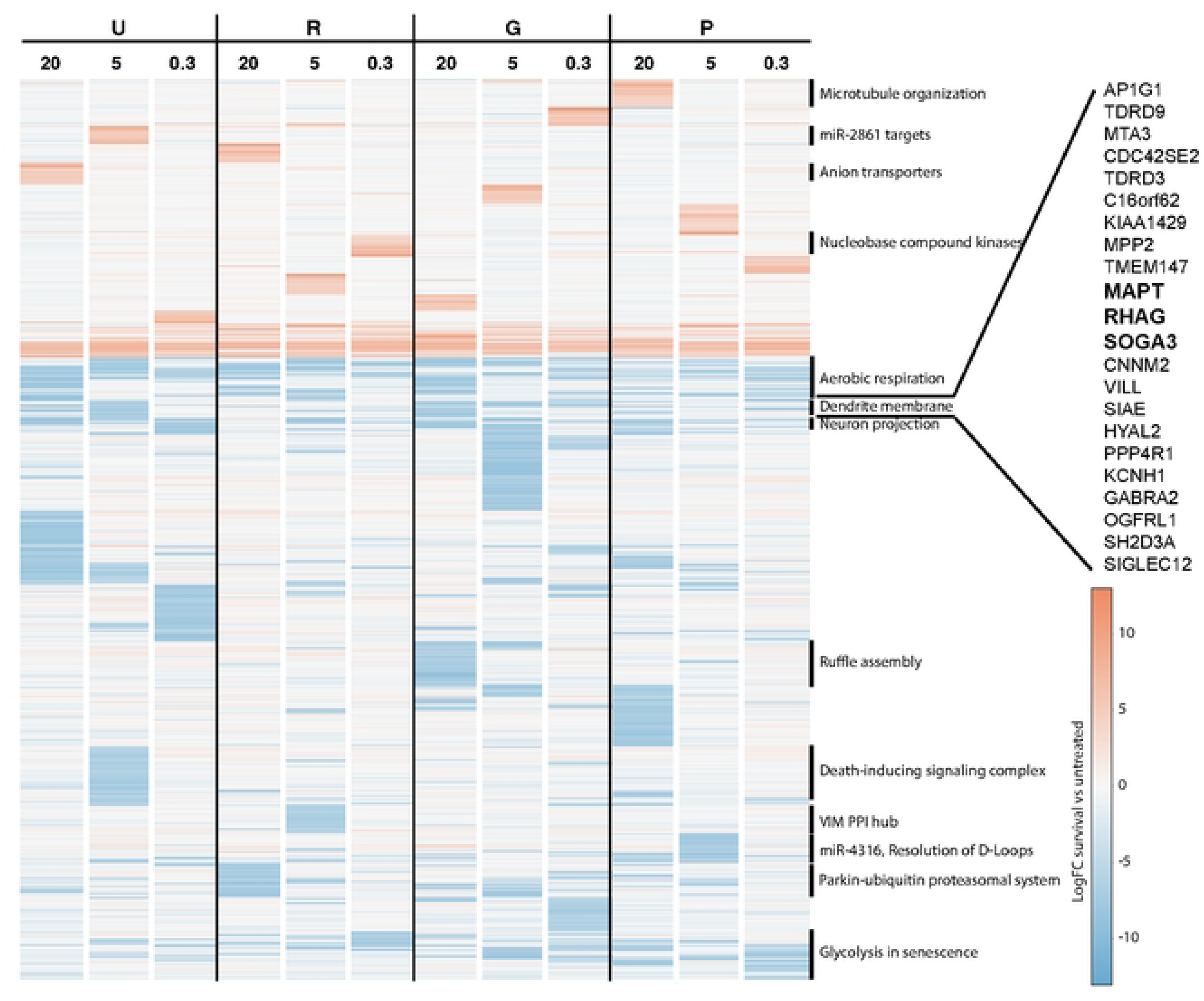
Hierarchical clustering identifies functionally coherent sets of neuronal metabolic survival genes. Genes with the strongest effect on neuronal metabolic survival in at least one metabolic state were clustered based on their average phenotype using the Ward D2 algorithm. Named gene sets were over-represented within functionally coherent clusters based on analysis with Enrichr, with adjusted p < 0.05, and more extensively catalogued in Table S5.

This approach revealed clusters of genes whose essentiality followed similar patterns of metabolic dependency. For instance, some genes were essential across all 12 conditions of the screen; this category was enriched for aerobic respiration genes (NDUFA8, NDUFA9, DLST, UQCRC2, adjusted p = 0.037, Fig 4). On the other hand, GO terms for the dendrite membrane and for neuron projections were enriched within sets of genes that were essential under high-oxygen glycolytic conditions. This observation further supports a functional connection in neurons between components of the neuron projection or dendrites and glycolytic function, and suggests that for specific genes, this connection may be sensitive to oxygen levels.

### Neuronal adaptation to reliance on glycolysis involves mitochondrial TCA cycle reprogramming

Although we found multiple mitochondrial pathways were essential for neuronal survival in the condition of dependence on respiration-only and high-oxygen conditions (Figs 2A, 2B, 2D, 3A, 4), genes essential for neuronal survival in high-oxygen glycolytic conditions were not as clearly annotated. In that setting, we were surprised to observe that several components of the TCA cycle were essential, especially MDH2 and DLST, which act in the later part of the cycle (Fig 5A). In contrast, pyruvate conversion to acetyl-CoA and earlier steps were more essential in high-oxygen respiratory conditions. TCA cycle genes were generally less essential for respiration-only or glycolytic-only survival at low-oxygen conditions (Fig 5B). These results indicate that altered TCA cycle activity is an important part of neuronal adaptation to a high-oxygen glycolytic state.

**Fig 5.**
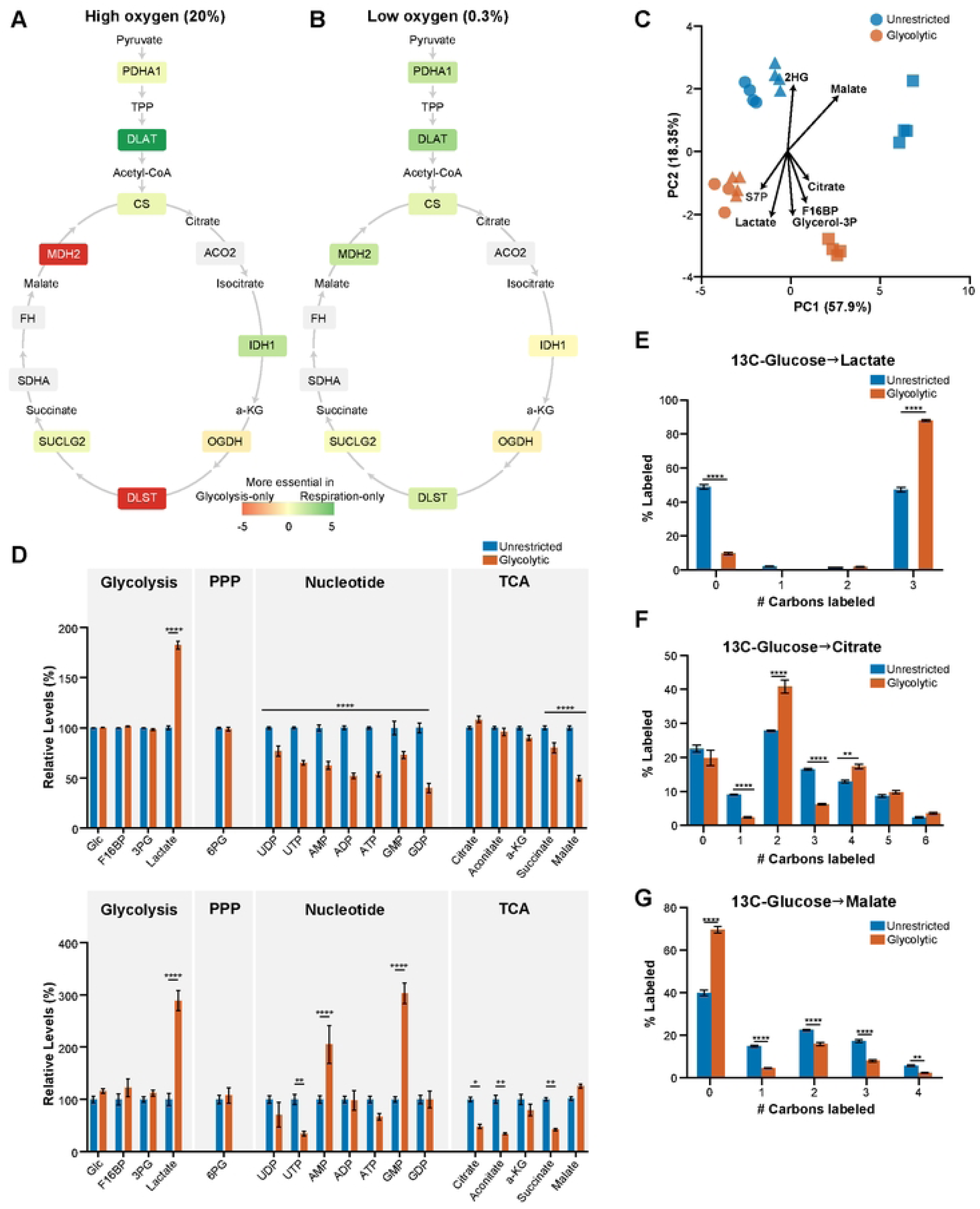
Neuronal survival in the high-oxygen glycolytic state is associated with metabolic reprogramming. A) Genes within the tricarboxylic acid (TCA) cycle have varied levels of essentiality under glycolytic versus respiratory conditions at high oxygen. B) TCA cycle genes are less essential for neuronal survival in hypoxic glycolytic and hypoxic respiratory conditions. C) Principal component analysis of 13C-glucose-derived labeling reveals metabolites that contributed to variation between high oxygen unrestricted (blue) and glycolytic metabolic states (red), including malate and citrate. Data compiled from n = 11 samples per glycolytic condition, n = 12 samples per unrestricted condition samples across three biological replicates (each shape corresponds to a separate biological replicate). D) Fraction of select 13C-glucose-labeled (top) and total amount of the same metabolites (bottom) under unrestricted (black) and glycolytic metabolic states (red) and high oxygen. E–G) Distribution of lactate (E), citrate (F) and malate (G) with zero, one or more carbons labelled by 13C-glucose under high-oxygen unrestricted and high-oxygen glycolytic conditions. Neuronal cells forced to depend on high-oxygen glycolysis incorporate more carbons derived from 13C-glucose into lactate and citrate pools, but fewer into malate pools. From n = 11 samples per glycolytic condition, n = 12 samples per unrestricted condition across three biological replicates for labeling experiments, n = 7 samples per glycolytic condition, n = 8 samples per unrestricted condition across two biological replicates for total metabolite pool size measurements. *p < 0.05, **p < 0.01, ***p < 0.001, ****p < 0.0001 by 2-way ANOVA with Šídák’s multiple comparisons test.

To confirm these observations, we performed targeted metabolomics in neurons expressing a non-targeting sgRNA. We found that labeled malate and citrate were major contributors to the differences between unrestricted and glycolysis-only states (Fig 5C). In glycolysis-only states, glucose carbons were used more for lactate production, and there were increased levels of low-energy nucleotide species (Fig 5D, 5E). This shift accompanied diversion of glucose carbons away from specific TCA cycle metabolites (succinate, malate). These changes in TCA cycle metabolites didn’t occur in respiration-only conditions with 13C-labeled pyruvate (Fig S1D). In high-oxygen glycolytic conditions, neurons increased glucose-derived carbon in citrate pools (Fig 5F) but decreased it in malate pools (Fig 5G). These findings suggest citrate-derived products are essential, while excess mitochondrial malate is harmful for neurons relying on glycolysis with abundant oxygen.

### Tau-encoding MAPT is a critical regulator of neuronal glycolysis

We were surprised to find *MAPT* among the genes that were essential for survival under glycolysis-only, high-oxygen conditions (Fig 6A, S2A). *MAPT* encodes tau, a protein that aggregates in tauopathies including Alzheimer’s disease [34] and FTD [35], as well as other neurodegenerative diseases such as Parkinson’s disease [36]. Much attention has been devoted to the toxicity of aggregated tau [37], and recent studies have proposed roles for tau in regulating glycogen metabolism [38] or supporting oxidative phosphorylation [39], although a direct role for tau in regulating glycolysis has not been characterized. Across the genome-wide CRISPRi screen, principal component analysis indicated that *MAPT* knockdown strongly influenced survival across glycolytic states (Fig S2B). The effect of MAPT knockdown was confirmed in multiple screens (Fig 6B, S2A, Table S6) and using multiple independent sgRNAs. In addition, relative to controls, neurons expressing CRISPRi sgRNA knocking down *MAPT* expression (Fig S3A) had decreased survival when cultured in glycolytic versus unrestricted conditions (Fig 6C, 6D).

**Fig 6.**
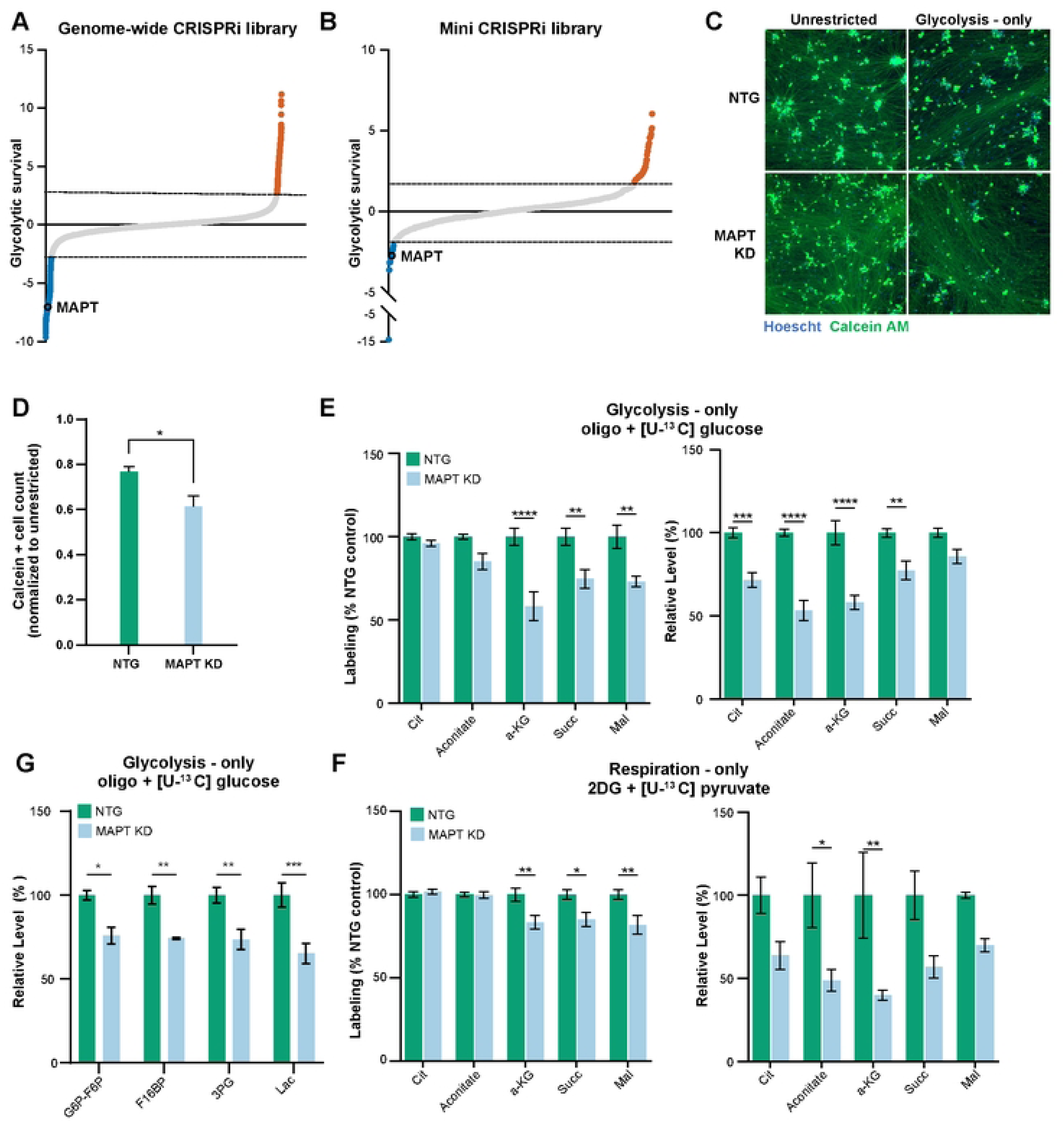
Tau regulates neuronal glycolysis. A, B) *MAPT* knockdown was associated with decreased neuronal survival in high-oxygen glycolytic conditions when targeted as part of a genome-wide dual-guide CRISPRi library (A) or a targeted single-guide mini CRISPRi library (B). Y-axis denotes the log-fold change in enrichment of the genes’ targeting sgRNAs after exposure to metabolic conditions relative to pre-exposure. Dashed lines mark 2 standard deviations from the average of non-targeting guides in each screen. Data is compiled from n = 2 replicates per metabolic state for the genome-wide screen, n = 4 replicates for unrestricted and untreated metabolic samples, n = 6 replicates for respiratory, physiologic, and glycolytic metabolic state for the mini-library screen. C-D) *MAPT* knockdown significantly decreases calcein-positive (green) neuronal cells counts in the high-oxygen glycolytic state. Cell counts were normalized to cell numbers in unrestricted high-oxygen conditions. Cells transfected with non-targeting (NTG) sgRNA serve as negative controls. Data compiled from n = 3 replicates. *p < 0.05 by 2-way ANOVA with Šídák’s multiple comparisons test. E,F) TCA cycle alterations in 13C-labeled (top graphs) and total (bottom graphs) metabolite pool sizes with *MAPT* knockdown were remarkably similar in glycolysis-only (E) or respiration-only (F). Data compiled from n = 4 samples. *p < 0.05, **p < 0.01, ***p < 0.001, ****p < 0.0001 by 2-way ANOVA with Šídák’s multiple comparisons test.

To understand if tau regulates glycolysis in neurons, we looked for clues among the genes with highest functional similarity to tau across the metabolic conditions. Knockdown of *RHAG* and *SOGA3* were the closest phenocopies to loss of *MAPT* genome-wide (Fig 4, Table S5). *RHAG* has been primarily studied in red blood cells, where it encodes a CO_2_ channel[40]. CO_2_ is a major waste product in respiration as a by-product of the TCA cycle. Therefore, we hypothesized that MAPT knockdown would impair CO_2_-generating steps within the TCA cycle. Indeed, targeted metabolomics confirmed that *MAPT* knockdown decreased glucose-derived carbon utilization into TCA cycle metabolite pools specifically at CO_2_-generating steps (generating α-ketoglutarate and succinyl-CoA), and also decreased TCA cycle metabolite pool sizes for these metabolites (Fig 6E). However, these same changes were observed in respiration-only conditions (Fig 6F), which were not observed to consistently result in a survival deficit in the genome-wide CRISPRi screen. Therefore, while our observations confirm that loss of *MAPT* expression decreases TCA cycle activity, this observation alone does not explain the survival deficit in the high-oxygen glycolytic state.

Another gene functionally similar gene to *MAPT*, *SOGA3*, also known as *MTCL3,* is known to regulate cytoskeletal cross-linking and to oppose gluconeogenesis, or reverse flow in glycolysis[41]. Notably, CO_2_ accumulation, a likely result from blocking CO_2_ channels, can cause reverse flux in the TCA cycle[42]. These observations led us to hypothesize that *MAPT* expression similarly opposes anabolic flow and promotes catabolic, forward flow of glycolysis. Using targeted metabolomics, we determined that neuronal cells with *MAPT* knockdown did indeed have smaller total pools of glycolytic metabolites, including lactate (Fig 6G). Notably, the carbon utilization within each glycolytic metabolite pool were mostly unchanged in neuronal cells with *MAPT* knockdown versus controls (Fig S2C). It is possible that during the 24 h incubation with 13C-glucose, the glycolytic pools became saturated with labeling, limiting the sensitivity to shifts in carbon utilization associated with *MAPT* expression.

To investigate if tau regulates glycolytic activity, we first queried an existing perturb-seq data set in which *MAPT* knockdown was performed in the same human iPSC-derived neuronal cell system we used[43]. In this data set, no individual glycolytic gene’s expression was significantly altered by *MAPT* knockdown (Fig S2D).

### The mRNA methyl transferase KIAA1429 regulates neuronal glycolytic gene expression

Another gene functionally similar to *MAPT* that was specifically required for neuronal survival in the high-oxygen glycolytic state, but not in unrestricted or respiratory conditions, was *KIAA1429*, which encodes the largest component of the mRNA methylation complex[44]. Due to this known role of *KIAA1429*, we hypothesized that *KIAA1429* selectively regulated expression of glycolytic enzymes, and might serve as an important regulator for the balance between respiration and glycolysis in neurons [45, 46]. Similar to our approach with *MAPT*, we first examined the perturb-seq dataset [32]. While sgRNAs targeting *KIAA1429* itself were not included in the experiment, we identified two CRISPR perturbations that were associated with decreased expression of *KIAA1429* (PPP2R2B knockdown p = 0.0011, FGD4 upregulation p = 0.021). Remarkably, the aerobic glycolysis pathway was enriched among the most downregulated genes across the average transcriptome when *KIAA1429* expression was low (Fig S4A). In contrast, mitochondrial and respiration-related pathways were not significantly represented among downregulated genes in cells with low *KIAA1429* expression. We expressed sgRNA targeting *KIAA1429* to knock down *KIAA1429* expression in neuronal cells (Fig S3B), and used RNAseq to confirm that the glycolytic pathway was significantly over-represented among downregulated genes in those cells (Fig 7A, S4B). The most downregulated glycolytic gene was *ALDOC,* which encodes an isoform of aldolase notably expressed in the hippocampus[47, 48].

**Fig 7.**
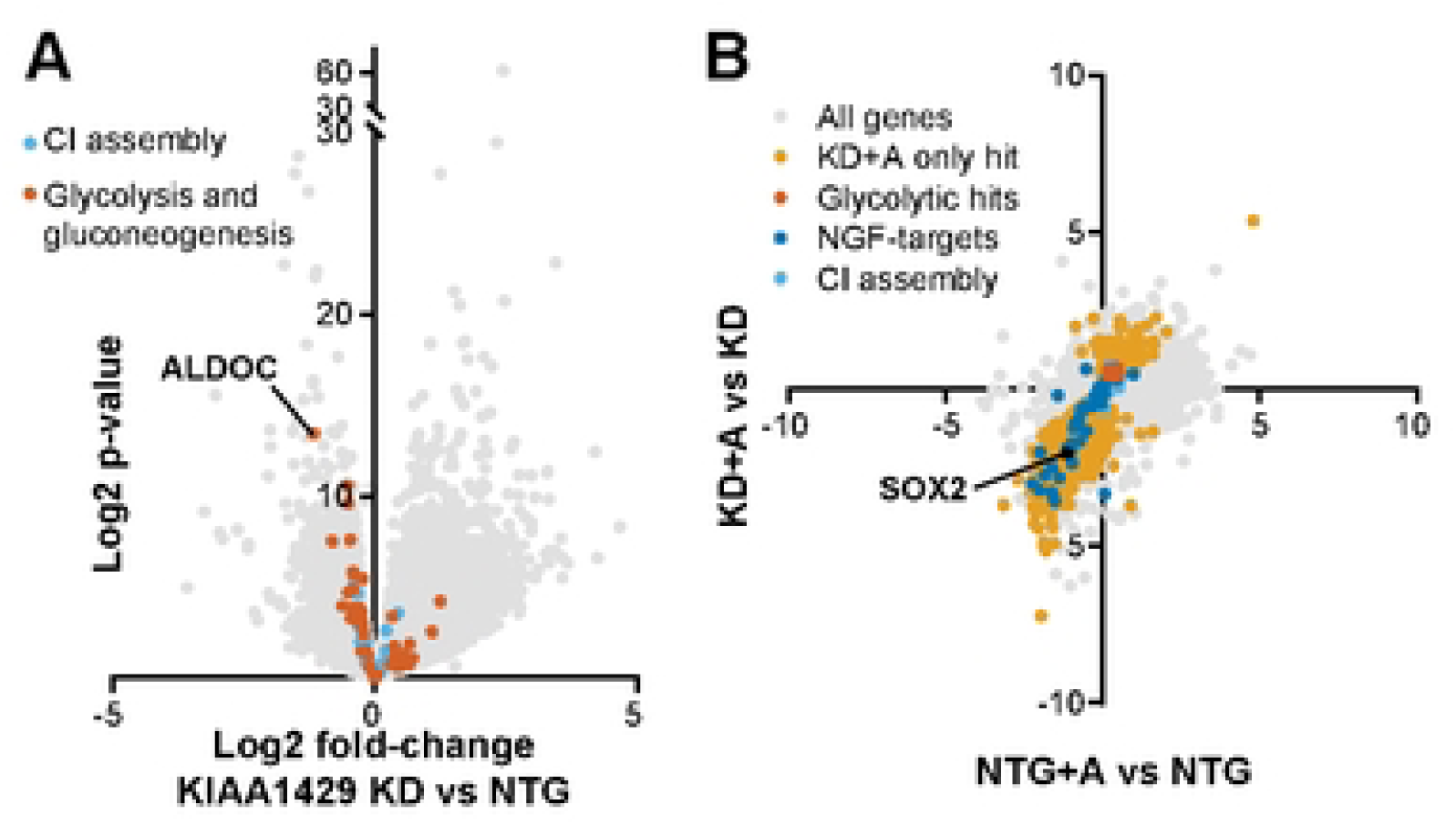
*KIAA1429* regulates glycolytic transcript abundance independent of mRNA stability. A) Volcano plot of genes up- or down-regulated in *KIAA1429* knockdown vs NTG control neurons. Glycolytic genes are more strongly affected than mitochondrial complex I assembly genes. Data compiled from n = 2 samples per condition. B) Plot of transcripts’ abundance in *KIAA1429* KD cells and NTG control cells after 4-hour exposure to transcription inhibitor actinomycin relative to before exposure. Knockdown in *KIAA1429* was associated with decreased abundance of *SOX2* and NGF-related transcription targets (dark blue) after treatment with actinomycin. The expression of these transcripts was not significantly decreased in neuronal cells expressing a non-targeting guide, also subjected to treatment with actinomycin, indicating that KIAA1429 activity has a stabilizing effect on these transcripts. In contrast, glycolytic transcripts (red) significantly increased expression in neuronal cells with *KIAA1429* knockdown and actinomycin treatment, indicating that stability of these transcripts was not impacted by *KIAA1429* expression. Data compiled from n = 2 samples per condition. All “hit” genes have adjusted p-values < 0.05.

mRNA transcripts can either be stabilized or degraded as a result of methylation[49–51], depending on the activity of interacting protein partners. To determine if knockdown of *KIAA1429* was associated with altered stability of glycolytic transcripts, we measured transcript abundance before and after a 4-hour treatment with actinomycin, a transcription inhibitor[52] (Fig 7B). Transcripts for which methylation has a stabilizing effect would be expected to decrease in abundance after actinomycin treatment in cells where *KIAA1429* is knocked down. Indeed, *SOX2*, a known target of stabilizing mRNA methylation[50], was significantly downregulated after actinomycin treatment in the cells with *KIAA1429* knockdown (Fig 7B). We also observed that many targets of NGF-stimulated transcription depend on *KIAA1429* expression for stability. While the culture media did not contain NGF, the overrepresentation of this particular annotated Reactome pathway may indicate that there is decreased sensitivity to pro-survival signaling from neurotrophins BDNF or NT-3, which were present in media. We did not observe a significant decline in glycolytic transcript abundance after inhibiting transcription with actinomycin. We conclude that *KIAA1429*-mediated transcript methylation does not stabilize glycolytic transcripts in neurons.

### Loss of functionally similar genes supporting neuronal glycolytic metabolism leads to elevated nucleotide synthesis

Our observations are consistent with a role for both *MAPT* and *KIAA1429* in supporting multiple steps in glycolysis in aerobic conditions. We hypothesized that loss of these and functionally similar genes would result in accumulation of early glycolytic metabolites or biosynthetic precursors derived from early glycolytic metabolites. To test this hypothesis, we created iPSC lines with targeted knockdown of *MAPT, KIAA1429,* and *TDRD3* (Fig S3), another gene whose knockdown similarly compromised survival in glycolytic conditions (Fig 8A). We also created human neurons with knockdown of *GLO1*, which also was essential in 20% O2-glycolytic conditions, but was functionally distinct from the other three genes (Fig S3).

**Fig 8.**
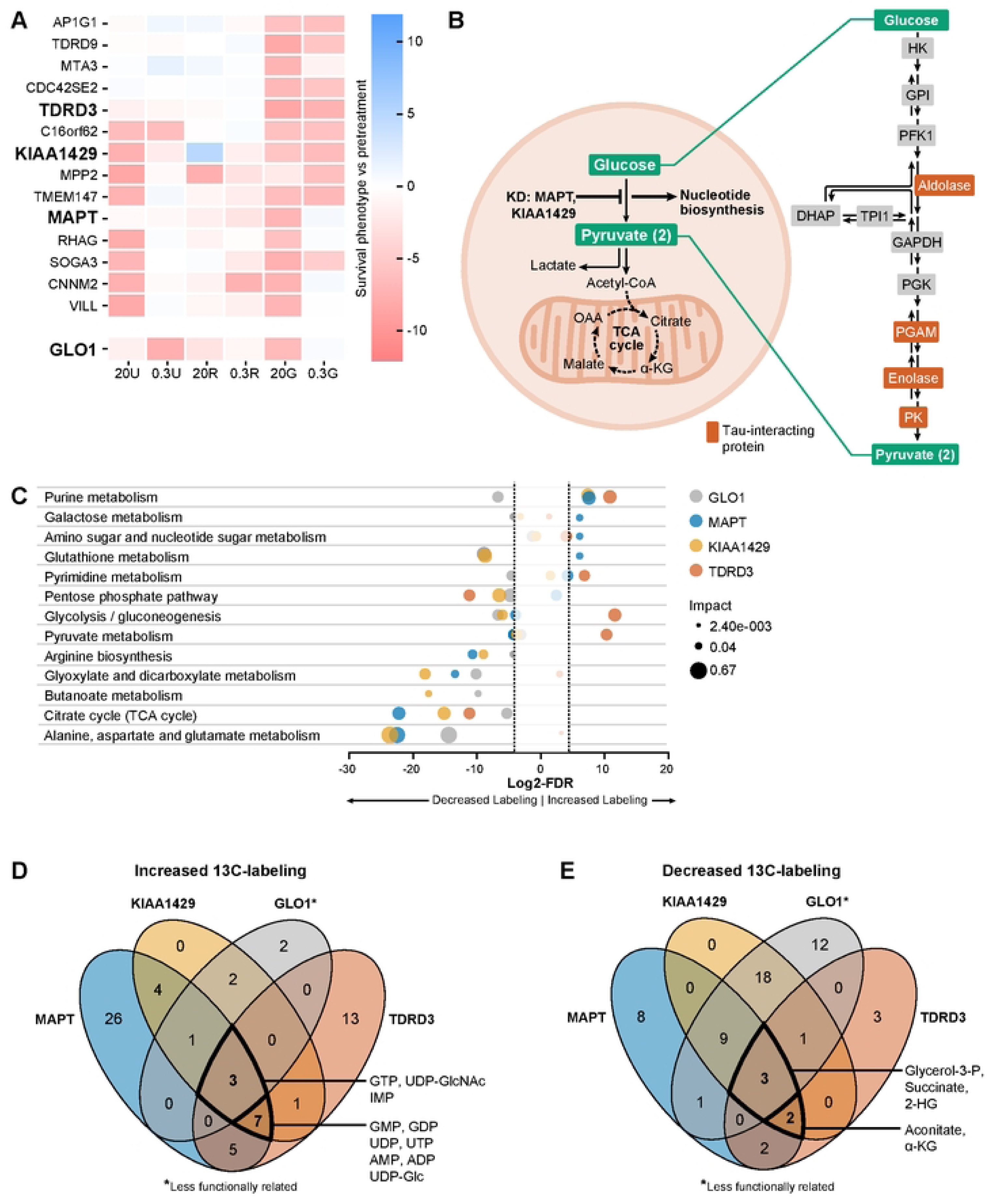
Diversion of carbon into purine pools is a defining feature of neurons with impaired survival in the high-oxygen glycolytic state. A) Heatmap of the cluster of genes enriched for dendrite membrane components from Fig 4 includes *MAPT*, *KIAA1429,* and *TDRD3*. Knockdown of these genes results in a common phenotype of poor survival in the high-oxygen glycolytic state (20G), while retaining survival in the high oxygen respiratory state (20R). B) Proposed mechanism by which knocking down genes that support various steps in late glycolysis results in diversion of glucose to upstream glycolytic branch metabolites, including nucleotides. C) Bubble plot of enriched metabolic pathways among metabolites with increased or decreased 13C-glucose derived labeling in neurons with genetic knockdowns. Bubble radius corresponds to pathway impact, which is a measure of the centrality of enriched metabolites. In the high-oxygen glycolytic state, knockdown of these three functionally similar genes in neuronal cells was associated with increased 13C-glucose-derived labeling across purine metabolites, compared to neuronal cells expressing a non-targeting sgRNA. In contrast, knockdown of *GLO1*, a gene less related gene by hierarchical clustering, did not increase 13C-glucose-derived labeling in purine metabolites. D) UDP, AMP, and ADP metabolites specifically are significantly increased 13C-glucose-derived labeling associated with knockdown in *MAPT*, *KIAA1429*, and *TDRD3* but not *GLO1* in the high-oxygen glycolytic state. E) TCA cycle metabolites are among the metabolites that have less 13C-glucose-derived labeling in the high-oxygen glycolytic state.

Unexpectedly, knockdown of *MAPT*, *KIAA1429*, and *TDRD3* all increased 13C-glucose-derived labeling of purine metabolite pools (Fig 8B, 8C, 8D), and decreased labeling of TCA cycle metabolite pools (Fig 8B, 8C, 8E). In contrast, knockdown of *GLO1* was not associated with elevated 13C-glucose labeling of purine metabolite pools. Specifically, AMP, ADP, and UDP had significantly more 13C-labeled carbons derived from glucose in human neuronal cells with *MAPT*, *KIAA1429*, or *TDRD3* knockdown (Fig 8D S5). Knockdown of *GLO1* was not associated with elevated 13C-labeled carbons in AMP metabolite pools, and was associated with decreased labeling in ADP and UDP metabolite pools. These observations support the hypothesis that *MAPT*, *KIAA1429*, and *TDRD3* support multiple steps within the later glycolytic pathway, and are consistent with excess nucleotide production being an important consequence of disrupted glycolysis in neurons.

## DISCUSSION

In this gene-by-environment screen for regulators of human neuronal survival across metabolic states, we highlight the systems-level importance of mitochondrial genes for survival when mitochondrial respiration is the sole source of cellular energy. Many of these genes are associated with known mitochondrial disorders, including Leigh syndrome. We hypothesize that neurodegeneration observed with these diseases could be modeled by combining genetic knockdown with metabolic conditions that force cells to rely on respiration. Furthermore, our work builds on our previous observations[13, 53, 54] that metabolic vulnerabilities caused by deficiencies in genes associated with Leigh syndrome could be ameliorated by therapeutic strategies that promote glycolysis, including hypoxia, by extending these findings into human neurons.

While it is well-known that neurons rely heavily on mitochondrial respiration for energy, neurons also require glycolysis for physiologic function[6, 7]. We induced a glycolysis-dependent, aerobic state in our screen by restricting mitochondrial respiration and forcing reliance on glycolysis for cellular energy in the presence of 20% O_2_. Not only does this approach increase sensitivity to regulators of glycolysis, it also models disease states where mitochondrial respiration is impaired, as well as neurodegenerative diseases[55, 56] and potentially aging[57]. Our work nominates a novel set of genes that are essential for neuronal survival in the high-oxygen glycolytic metabolic state. Indeed, we observed that neuronal cells adapting to reliance on glycolysis in high oxygen conditions not only increased lactate production from glucose, but also remodel TCA cycle activity. This TCA remodeling adaptive process involved using less carbon from glucose in TCA metabolite pools, which is consistent with models of altered TCA flow observed in cellular state change, where those deficits could be explained by diversion of carbons towards alternative metabolic pathways, or by the catabolism of alternative carbon sources [58, 59]. Taken together, these metabolic reprogramming steps could be understood as a shift away from catabolic energy harnessing towards anabolic carbon and energy conservation. While our proposed model focuses on genes that regulate glycolytic function, we do not exclude the possibility that a shift in TCA cycle activity towards a more carbon-conserving and remodeled state are also important parts of supporting neuronal glycolytic function.

We observe that deficiencies in genes associated with autism shift developing neuronal cells away from reliance on mitochondrial respiration, and towards a more flexible metabolism that promotes survival in glycolytic and hypoxic conditions. These observations are consistent with work by several groups[60–62]. Our data furthermore show that loss of function in multiple ASD-related genes causes convergent shifts in dependence on specific metabolic substrates. Notably, knockdown of genes associated with ASD did not behave similarly across all metabolic conditions. Further studies should illuminate which metabolic factors have the greatest influence on these ASD-associated genes, as well as how ASD-related proteins interact with metabolic energy pathways or energy-regulating transcription factors, particularly those that are active when glucose is abundant and oxygen scarce. This interaction could serve as a target for metabolic therapy in the treatment of ASD.

One important regulator of neuronal survival in the high-oxygen glycolytic state is *KIAA1429*, a major component of the RNA methylation complex. We used perturb-seq datasets to predict what glycolytic gene expression would be co-regulated with *KIAA1429*. This work demonstrates the value of perturb-seq datasets in dissecting gene circuits, and specifically in understanding epitranscriptome regulatory mechanisms, including RNA methylation. We confirmed that *KIAA1429* expression was critical to maintain expression of glycolytic transcripts, particularly *ALDOC*, however, expression of *KIAA1429* did not impact glycolytic transcript stability. RNA methylation has been observed to regulate transcript translation efficiency[63] and transport within cells[64, 65]. Further investigation will be required to determine whether appropriate localization and translation of glycolytic transcripts within dendrites is a key determinant of neuronal glycolytic function and survival in the high-oxygen glycolytic state.

We found that *MAPT*, which encodes tau, was essential for neuronal survival in the high-oxygen glycolytic metabolic state. Tau forms aggregates in a wide range of neurodegenerative disease, and *MAPT* mutations are associated with dementia. However, little is understood about the physiologic functions of tau, particularly in regulating metabolic processes. Tracy et al. found that dementia-associated mutations reduce tau interaction with mitochondrial proteins and impair mitochondrial bioenergetics[66]. We find that loss of MAPT expression is not associated with a decrease in survival when neuronal cells are forced to rely on respiration. Rather, we observed a decrease in survival specifically when neuronal cells are forced to rely on high-oxygen glycolysis. [67] Notably, Tracy et al.’s dataset of tau protein interactors includes several glycolytic enzymes, particularly among the later glycolytic steps. Our observations and this dataset suggest a potential role for tau in organizing glycolytic activity through coordinated interaction with multiple glycolytic enzyme partners. Assemblies of glycolytic enzymes, sometimes referred to as a glycolytic metabolon, have been observed in association with the cytoskeleton[68] and on the surface of mitochondria[69]. It is possible that tau plays an essential role in glycolytic coordination and catalysis in these cellular compartments. Loss of glycolytic-supporting functions of tau, which might occur as a result of aging-associated aggregation of disordered proteins[70, 71], could lead to energy failure in neurons that have already begun to shift towards dependence on glycolysis[67]. Not only does our work highlight the need to understand the essential role tau plays in regulating glycolysis, it also indicates that some toxicity associated with loss of tau could be ameliorated by bypassing glycolysis and facilitating mitochondrial respiration.

Finally, we observed that a shared feature of neuronal cells that were deficient in high-oxygen glycolytic-essential genes was increased diversion of carbons from glucose into nucleotide pools. In the case of *KIAA1429* and *MAPT* knockdown, we assert that this effect is mediated by coordinated loss of function at multiple glycolytic steps, likely causing a diversion of glucose-derived carbons in the early glycolytic steps into the pentose phosphate pathway shunt. It remains unclear whether the metabolic shifts observed in neuronal cells adjusting to the high-oxygen glycolytic state—such as altered TCA cycle activity, increased lactate production, or reduced nucleotide synthesis—are adaptive responses, and to what extent each contributes to neuronal survival. Notably, these genes that are essential in the high-oxygen glycolytic state are not essential when neuronal cells are forced to rely on respiration. This suggests that loss of these genes does not directly impair mitochondrial function. Rather, loss of these genes combined with inhibition of mitochondrial respiration leads to toxicity. We have previously observed that the decision point between carbon utilization in the catalytic glycolytic pathway versus the anabolic pentose phosphate pathway can contribute to energy failure by exacerbating energy consumption[13]. We provide further support that managing this balance through pathway-level support of glycolysis regulates neuronal survival in the high-oxygen glycolytic state. Further studies will be needed to better understand the molecular mechanisms that underlie this regulation, and to understand if supporting mitochondrial respiration is sufficient to promote neuronal survival when glycolysis fails.

## METHODS

### Cell Culture

Human iPSCs in this study (male WTC11 background) were provided by Martin Kampman’s lab. These cells have stable integration of dCas9-KRAB (CRISPRi) and an inducible neurogenin-2 (NGN2) expression system received from Li Gan’s lab[10]. iPSCs were maintained at 37 °C and cultured in Gibco StemFlex Medium (Cat. No. A3349401) on cell culture flasks (Corning; assorted Cat. No.) coated with Matrigel hESC-Qualified Matrix, LDEV-free (Corning; Cat. No. 354277) diluted 1:121 DMEM F12 (GIBCO/Thermo Fisher Scientific; Cat. No. 11320033). Briefly, StemFlex Medium was replaced every other day. When 80-90% confluent, cells were passaged by aspirating media, incubating with Accutase (STEMCELL Technologies; Cat. No. 07920) at 37 °C for 5 min., diluting Accutase 1:5 in DPBS, collecting in conicals, centrifuging at 300 g for 5 min, aspirating supernatant, resuspending in StemFlex Medium supplemented with .01mM Y-27632 dihydrochloride ROCK inhibitor (Tocris; Cat. No. 125410), counting, and plating onto Matrigel-coated plates at desired number.

To differentiate iPSCs into neurons, iPSCs were released and centrifuged as above, and pelleted cells were resuspended in N2 Pre-Differentiation Medium containing the following: Knockout DMEM/F12 (GIBCO/Thermo Fisher Scientific; Cat. No. 12660-012) as the base, 1X MEM Non-Essential Amino Acids (GIBCO/Thermo Fisher Scientific; Cat. No. 11140-050), 1X N2 Supplement (GIBCO/Thermo Fisher Scientific; Cat. No. 17502-048), 10ng/mL NT-3 (STEMCELL Technologies; Cat. No. 78074), 10ng/mL BDNF (STEMCELL Technologies Cat. No. 78005.1), 1 µg/mL Mouse Laminin (Thermo Fisher Scientific; Cat. No. 23017-015), 10nM ROCK inhibitor, and 2 µg/mL doxycycline hyclate (Sigma-Aldrich; Cat. No. D9891) to induce expression of mNGN2. iPSCs were counted and plated onto Matrigel-coated plates in N2 Pre-Differentiation Medium for three days. On day three (Day -1), the cells were fed with N2 Pre-Differentiation Medium containing doxycycline hyclate at a 1:1000 dilution.

After three days (Day 0), pre-differentiated cells were released, centrifuged as described above, and the pelleted cells were resuspended in Classic N2/B27 Neuronal Medium containing the following: half DMEM/F12 (GIBCO/Thermo Fisher Scientific; Cat. No. 11320-033) and half Neurobasal-A (GIBCO/Thermo Fisher Scientific; Cat. No. 10888-022) as the base, 1X MEM Non-Essential Amino Acids, 0.5X GlutaMAX Supplement (GIBCO/Thermo Fisher Scientific; Cat. No. 35050-061), 0.5X N2 Supplement, 10 ng/mL NT-3, 10 ng/mL BDNF, 1 µg/mL Mouse Laminin, and 2 µg/mL doxycycline hyclate.

Pre-differentiated cells were counted and plated at the desired density onto plates coated with Poly-L-Ornithine solution (Sigma-Aldrich; Cat. No. P4957-50ML) in Classic N2/B27 Neuronal Medium. On day 2, the media was replaced with fresh Classic N2/B27 Neuronal Medium without doxycycline. On day 7 and every 6 days thereafter, half of the medium was replaced with an equal volume of fresh Classic N2/B27 Neuronal Medium without doxycycline. For experiments involving metabolic media treatments, all media was removed on day 18 and replaced with metabolic media, as described below. All experiments were completed on day 21 post-differentiation.

### CRISPRi gene knockdown in human neurons

Lentivirus expressing the dual-sgRNA[8] or single-guide sgRNA[13, 72] libraries were produced by Vectorbuilder and the UCSF Viracore. This mini-library was enriched in sgRNAs targeting genes that are essential for mitochondrial ATP[53], and expanded to include genes associated with Alzheimer’s or Parkinson’s disease (Table S6). Notably, this mini-library used single-guide gene targeting[72] and different sgRNA targeting sequences, making validated phenotypes unlikely to be caused by an artifact of the particular sgRNA used. Individual dual-sgRNA constructs targeting specific genes were generated by express mutagenesis (Genscript) from a single dual-sgRNA construct supplied by Jonathan Weissman’s lab. Lentivirus expressing individual dual-sgRNA constructs were generated as described by Tian et al[9]. HEK293T cells were plated to 80% confluence in 6-well plates. We then diluted 1μg transfer plasmid and 1μg of third generation packaging mix into 200 μL OPTIMEM and 6 μL of TransIT-Lenti Reagent. This transfection mix was then incubated for 10 minutes before addition to the HEK media. After 48 hours, supernatant was collected and filtered through a syringe equipped with a 0.45 μm PVDF filter. Lentivirus precipitation solution (Alstem) was added to this filtrate and stored at 4 °C for 24 hours, before centrifugation for 30 minutes at 1500xg in a centrifuge pre-chilled to 4 °C. The virus-containing pellet was then resuspended into StemFlex media and flash frozen in liquid nitrogen.

CRISPRi sgRNA were transduced with polybrene (8 μg/mL) into iPSCs. Four days after transduction, cells were expanded and cultured with 1 μg/mL puromycin for 5 days to select for cells expressing CRISPRi sgRNA constructs. iPSCs expressing CRISPRi sgRNA were differentiated into human neuronal cells as described above.

### Metabolic media treatments

For metabolic media treatments, neuronal cells were cultured for 3 days in either unrestricted media (21.25 mM glucose, 0.36 mM pyruvate, 2.25 mM glutamine, matching the substrate concentrations of N2/B27 neuronal differentiation media), respiration-only media (10 mM pyruvate, 10 mM 2-deoxy-D-glucose, 2.25 mM glutamine), glycolysis-only media (2 mM glucose, 5 μM oligomycin A, 2.25 mM glutamine), or physiologic media (1.5 mM glucose[73–75], 0.2 mM pyruvate[76], 0.2 mM β-hydroxybutyrate[21], 1 mM lactate[76–78], 0.75 mM glutamine[79], approximating the physiologic state based on literature values based primarily on microdialysis measurements). These media conditions were combined with either normal room levels of oxygen, normal brain levels of oxygen, or hypoxia (20%, 5%, and 0.3% O_2_).

### CRISPRi screening

Neuronal cells were dissociated with accutase and collected by centrifugation at 300xg for 5 minutes before and after treatment with metabolic medias. Genomic DNA was isolated using the Macherey-Nagel Nucleospin Tissue kit. The sgRNAs were amplified and adaptors attached in a single PCR step, with a total of no more than 4 μg of undigested genomic DNA used per 100 μL PCR reaction. For the dual-sgRNA CRISPRi screen, PCR was conducted using NEBNext Ultra II Q5 MasterMix using reverse primer: CAAGCAGAAGACGGCATACGAGATGCGGCCGGCTGTTTCCAGCTTAGCTCTTAAA and forward primer: AATGATACGGCGACCACCGAGATCTACACNNNNNNNNCGCGTATCCCTTGGAGAACCACC, which included sequencing adaptor and indexing sequences, and where *N* refers to a variable index sequence. For the single sgRNA library screen, PCR was conducted using Q5 HotStart High Fidelity Polymerase (NEB) using forward primer: aatgatacggcgaccaccgaGATCGGAAGAGCACACGTCTGAACTCCAGTCACNNNNNNgcacaaaaggaaactcaccct and reverse primer: caagcagaagacggcatacgaCGACTCGGTGCCACTTTTTC, where N refers to a variable index sequence. PCR parameters were 98 °C for 30 s, followed by 22 cycles of 98 °C for 10 s, 63 °C for 75 s, and ending with 72 °C for 5 min, followed by holding at 4 °C. Resulting PCR product from the multiple reactions were pooled, followed by removal of unincorporated primers and gDNA using the GeneRead Size Selection Kit. Quality, size, and purity of the PCR product was assessed by bioanalyzer (Agilent) before sequencing on an Illumina NovaSeq 6000 for the dual-sgRNA library, or an Illumina HiSeq 2500 for the single-sgRNA library. Custom primers were used for the dual-sgRNA sequencing strategy, including the read 1 primer (R1): CGCGTATCCCTTGGAGAACCACCTTGTTGG, index 1 primer (I1): TGCTCGAATCTACACTCAGCTATGGCGCTG, read 2 primer (R2): GCGGCCGGCTGTTTCCAGCTTAGCTCTTAAAC, for a paired-end 50 run with R1: 19, I1: 8, R2: 19. A given gene perturbation’s survival phenotype was calculated using the MAGeCK RRA algorithm[80] with total normalization method. We defined the strongest “hit” genes as those that were at least three standard deviations from the mean of the non-targeting controls in at least one metabolic condition, had an FDR < 0.05, and did not have zero counts in any sample prior to the metabolic stress.

### CRISPRi Screen Overrepresentation Analysis

Enrichr^5,6^ was used to perform enrichment analysis of gene lists. To generate the gene lists from CRISPRi screen phenotypes, the 500 genes with the highest log fold changes and the 500 genes with the lowest log fold changes were chosen. Gene lists were compared against the human gene sets KEGG 2021, Reactome 2022/2024, Wikipathways 2023/2024, GO biological processes 2023, GO cellular component 2023, GO molecular function 2023, ClinVar 2019, Orphanet Augmented 2021, and DisGeNET v24.1. Fisher’s exact test *P* values were adjusted for multiple testing using the Benjamini-Hochberg procedure^7^. For over-representation analysis of clusters of “hit” genes determined by Ward.D2 hierarchical clustering, we submitted clusters of genes into gProfiler or Enrichr and selected pathways with p-adj > 0.05.

### Network Analysis

For gene network analyses of “hit” genes, the STRING interaction database was used to reconstruct gene networks using stringApp for Cytoscape. Clustering was performed on the 1163 node network using ClusterMaker2^10^. Markov Clustering (MCL) was selected as the network partitioning algorithm to divide networks into densely connected subnetworks and RCy3 was used to facilitate finding a reasonable parameter for MCL granularity. Clusters with more than 10 genes are selected, and the clusters are rearranged according to the number of genes.

### Targeted metabolomics

Human iPSC-derived neurons were differentiated and matured for 19 days prior to undergoing 24h incubation in media containing [U-^13^C] metabolic probes. Metabolic media included varying concentrations of glucose and pyruvate in a base media composed of Neurobasal A without D-glucose or sodium pyruvate (Gibco A2477501), 1X NEAA (Gibco 11140050), 2.25mM GlutaMAX (Gibco 35050061), 0.5X N2 supplement (Gibco 17502048), 0.5X B27 supplement (Gibco 17504044), 10 ng/mL NT-3 (Stemcell Tech 78074), 10 ng/mL BDNF (Stemcell Tech 78005), and

1 μg/mL of laminin (Gibco 23017015). Each media condition was supplemented with [U-^13^C]glucose (Cambridge Isotope Labs CLM-1396–1) or [U-^13^C]pyruvate (Cambridge Isotope Labs CLM-2440-1) for targeted metabolomics. Unrestricted media conditions included 21.25 mM [U-^13^C]glucose, 2.5 mM pyruvate, glycolysis-only conditions included 21.25 mM [U-^13^C]glucose and 5 μM oligomycin A, respiration-only media included 2.5 mM [U-^13^C]pyruvate and 10 mM 2-deoxyglucose. After 24h incubation, metabolites were extracted via cellular wash in ammonium acetate at 4C (150 mM, pH 7.4), incubation in 80% methanol at -80C for 30 min, and centrifugation at 14,000 rpm for 10 min at 4C. Metabolite supernatants were dried in a Labconco CentriVap without heat, prior to storage at -80C. Metabolites were quantified by the UCLA Metabolomics Core.

### Transcriptomics Analysis

To identify transcriptionally perturbed genes and pathways regulated by KIAA1429, neuronal cells expressing either KIAA1429-targeting sgRNA or a non-targeting sgRNA were cultured for 21 days.

To identify transcripts whose stability was modified by KIAA1429 expression, we treated 21-day aged neuronal cells expressing either KIAA1429-targeting sgRNA or non-targeting sgRNA with 10 μg/mL actinomycin D for 4 hours[52]. We used accutase to dissociate cells and collected cell bodies by centrifugation at 300 x g. Cell pellets were flash frozen in liquid nitrogen before submitting to BGI. Total RNA was isolated with Qiagen RNeasy Mini Kit (Qiagen, Germany) according to the manufacturer’s protocol. Isolated RNA sample quality was assessed by High Sensitivity RNA Tapestation (Agilent Technologies Inc., California, USA) and quantified by AccuBlue® Broad Range RNA Quantitation assay (Biotium, California, USA). Ribosomal RNA depletion was performed with QIAseq® FastSelect -rRNA HMR kit (Qiagen, Hilden, Germany) per manufacturers’ instructions. All library construction were prepared according to the NEBNext® Ultra™ II Directional RNA Library Prep Kit for Illumina® (New England BioLabs Inc., Massachusetts, USA). Final libraries quantity was assessed by Qubit 2.0 (ThermoFisher, Massachusetts, USA) and quality was assessed by TapeStation D1000 ScreenTape (Agilent Technologies Inc., California, USA). Final library size was about 430bp with an insert size of about 300bp. Illumina® 8-nt dual-indices were used. Equimolar pooling of libraries was performed based on QC values and sequenced on DNBseq T7 platform (MGI Tech Co, Shenzhen, China) with a read length configuration of 150 PE for 60M PE reads per sample (30M in each direction). Raw sequencing data was filtered using SOAPnuke[81] for quality control. Filtered sequencing reads were analyzed in Galaxy[82]. HISAT2 was used to align reads against the homo sapiens hg38.v2201 reference genome[83]. The FeatureCounts tool was used to quantify read counts[84]. The DESeq2 tool was used to identify differentially expressed genes[85]. Enrichr was used to perform enrichment analysis of differentially expressed genes (P < 0.05).

To identify genetic pathways that were co-perturbed in iPSC-derived neuronal cells with low KIAA1429 expression, we first identified all sgRNA used in the Tian et. al. 2019[9] and Tian et. al. 2021[43] with decreased KIAA1429 expression (P-value < 0.05). We averaged the log-fold change in expression for each transcript across these perturbed transcriptomes. Enrichr was used as described above to perform enrichment analysis of the 250 genes with lowest expression in this aggregate transcriptome.

### Metabolic Pathway Enrichment Analysis

In order to determine which metabolic pathways were enriched among metabolite pools with alterations in labeled carbons derived from 13C-labeled metabolic substrates, we used the Pathway Enrichment tool in Metaboanalyst[86]. KEGG identification numbers for metabolites with increased or decreased labeling in knockdown lines relative to non-targeting controls were uploaded to Metaboanalyst, and enriched pathways were identified using the Hypergeometric Test. Pathway impact was quantified based on the relative-betweeness centrality of the metabolite within pathway network topology within the KEGG pathway library, updated as of December 2023. We selected pathways to display where at least one gene knockdown was associated with an FDR < 0.05. For pathways that included metabolites with increased and decreased labeling, we plotted the more extreme log-FDR and corresponding pathway impact.

### RNA Isolation, reverse transcription (RT), and real-time RT-qPCR

We used the RNeasy Mini Kit (Qiagen) to isolate total RNA from cell pellets of approximately 500,000 cells according to the manufacturer’s protocol. cDNA was synthesized with the High-Capacity RNA-to-cDNA Kit (ThermoFisher), using a manufacturer-optimized transcription cycle (37 °C for 60 min, inactivation at 95 °C for 5 min, hold at 4 °C). Gene expression was measured by real-time PCR on QuantStudio 5 using FAM-MGB Taqman Gene Expression Assays (ThermoFisher, assay ID: Hs00198702_m1 *GLO1*, Hs00936421_m1 *KIAA1429*, Hs01556304_m1 *TDRD3*, Hs00902194_m1 *MAPT*, Hs00175976_m1 *HK1*, Hs00976715_m1 *GPI*, Hs01075411_m1 *PFKM*, Hs01378790_g1 *LDHA*, Hs00761782_s1 *PKM*, Hs01652468_g1 *PGAM1*, Hs00361415_m1 *ENO1*) together with VIC-MGB human ACTB (ThermoFisher, #4326315E). RT-qPCR reactions were performed using standard PCR conditions (UDG incubation - 50 °C for 2 min, enzyme activation - 95 °C for 10 min, followed by PCR cycle – 95 °C for 15 s, 60 °C for 1 min, repeated for 40 cycles) in a 384-well plate, in duplicate and from 2-3 independent experiments. CT (threshold cycle) values of each gene were averaged and calculated relative to CT values of ACTB using the 2^-ΔΔCT^ method.

### Viability Assay

After 3 days of pre-differentiation, human iPSCs were replated into 96-well black plates with clear bottoms (Greiner), pre-coated with 75 µL of poly-L-ornithine (PLO) per well at room temperature for 2 days, followed by laminin (5 µg/mL) in a cold room for 1 day. Each well was seeded with 25,000 cells in 100 µL of neuronal medium. Medium was nearly completely replaced every 6 days, leaving a minimal volume to prevent wells from drying out. On day 18 of differentiation, metabolically restricted and control media were applied through a full medium change. To simulate screening conditions, only 50 µL of medium was added to each well during the 3-day treatment period. Cell viability was assessed by staining neurons with Hoechst and Calcein-AM. A 50 µL volume of 2× staining solution was added to each well and incubated for 15 minutes before imaging on the CX7 High-Content Screening System. Cells positive for both Hoechst and Calcein-AM signals were classified as live cells. Nine fields of view were captured per well to ensure a representative assessment of cell viability. The resulting images were hyperstacked using a custom-written Fiji macro and analyzed with CellProfiler[87].

## ACKNOWLEDGEMENTS

We thank Eric Chow and the UCSF Center for Advanced Technology, as well as Francoise Chanut for helping edit the manuscript, Tami Tolpa for help with graphics and Vy Ho for administrative assistance. This work was funded in part by R01 AG065428 (K.N., I.J.), the UCSF Bakar Aging Research Institute (BARI, K.N.), and the UCSF Nutrition Obesity Research Center (NORC). It was also supported by a Berkelhammer Award for Excellence in Neuroscience (N.B.), F32 AG063457-02 (N.B.) and K01AG078485 (N.B.). JXM was supported by the NIH/NIGMS Institutional Research and Academic Career Development Award (IRACDA) at UCSF (K12FM081266), and YSM and JY were supported by a Hillblom Center & BARI Graduate Fellowship. We thank the Gladstone Flow Cytometry Core for assistance and use of flow cytometer equipment, supported by NIH S10RR028962, NIH P30 AI027763, and the James B. Pendleton Charitable Trust, and Anke Meyer-Franke at the Gladstone Assay Development and Drug Discovery Core for assistance with high-content microscopy. We thank the UCLA Metabolomics Center for their assistance with stable isotope labeling metabolomics.

## Supplementary Figure Legends

Supplementary Fig 1. A) STRING-based physical interactions combined with survival screen phenotypes nominate important roles for β-arrestin (encoded by gene *ARBB1*) and specific olfactory receptors OR6K3 and OR5M10 in regulating neuronal survival in hypoxic conditions with unrestricted metabolism. Similarly, this analysis nominates activity of the SCF E3 ubiquitin ligase complex, which depends on NEDD8 and CDC34, as essential for neuron survival in hypoxic conditions, with either unrestricted metabolism or when metabolism is restricted to respiration-only. B) Knockdown of few glycolytic enzyme-encoding genes lead to decreased glycolytic survival when neuronal cells are forced to rely on glycolysis-only in high oxygen conditions. Glycolytic enzymes are colored based on their log-fold change in enrichment of their associated targeting sgRNA in the genome-wide CRISPRi screen post-shift in metabolic conditions versus pre-treatment. B) Knockdown of glycolytic enzyme-encoding genes also did not have strong effects on survival when neuronal cells are forced to rely on respiration-only. Data for genome-wide CRISPRi screens compiled from n= 2 screens. C) Neuronal cells forced to rely on respiration-only incorporated significantly more carbons derived from 13C-pyruvate into lactate and TCA cycle metabolite pools, compared to neuronal cells cultured in unrestricted metabolism conditions. However, neuronal cells forced to rely on respiration-only had total TCA metabolite pool sizes that were not significantly different than in unrestricted metabolism conditions, except that they had significantly smaller malate pool, increased total levels of purine metabolites with fewer high-energy phosphate bonds (AMP, GMP), and significantly smaller total levels of nucleotides with more high-energy phosphate bonds (UTP, ATP). *p < 0.05, **p < 0.01, ***p < 0.001, ****p < 0.0001 by 2-way ANOVA with Šídák’s multiple comparisons test.

Supplementary Fig 2. A) *MAPT* knockdown was not associated with decreased neuronal survival in unrestricted, physiologic, or respiratory survival when targeted as part of a genome-wide dual-guide CRISPRi library or a targeted single-guide mini CRISPRi library. Dashed lines depict 2 standard deviations from the average of non-targeting guides in each screen. Data is compiled from n = 2 replicates per metabolic state for the genome-wide screen, n = 4 replicates for unrestricted and untreated metabolic samples, n = 6 replicates for respiratory, physiologic, and glycolytic metabolic state for the mini-library screen. B) Loading for *MAPT* and *RRAD* highlight that the variation *MAPT* expression correlates with the variation across metabolic states examined in the genome-wide CRISPRi screen along the first principal component. Notably, the first principal component explains variation across glycolytic states across the range of oxygen levels. *RRAD* expression correlates with variation across metabolic states along the second principal component, which accounts for separation between glycolytic and non-glycolytic metabolic states. C) Neuronal cells with knockdown in *MAPT* expression have no significant change in 13C-glucose derived labeling in high-oxygen glycolytic conditions, compared to neuronal cells expressing a non-targeting sgRNA, with the exception of increased 13C-glucose derived labeling in the glucose-6-phosphate/fructose-6-phosphate metabolite pool. Data compiled from n = 4 samples. ****p < 0.0001 by 2-way ANOVA with Šídák’s multiple comparisons test. D) Neuronal cells expressing the *MAPT*-targeting CRISPRi sgRNA in the Tian et. al perturb-seq dataset[43] do not have significant alterations in expression of glycolytic transcripts.

Supplementary Fig 3. Individually created neuronal cells lines have robust CRISPRi-mediated knockdown of targeted genes *MAPT*, *KIAA1429*, *GLO1*, and *TDRD3*, measured by RT-qPCR. Data compiled from n = 2 independent replicates. ****p < 0.0001 by t-test with Welch’s correction.

Supplementary Fig 4. A) Aerobic glycolysis-related genes feature among the top 5 pathways in Human Wikipathways database, based on enrichment score, which were over-represented among the most co-downregulated genes across CRISPR-perturbations with significantly downregulated *KIAA1429* expression, compiled from the Tian et. al. 2019 and Tian et. al. 2021 perturb-seq datasets[9, 43]. B) Aerobic glycolysis-related genes also feature among the top 5 pathways in Human Wikipathways database, based on enrichment score, which were over-represented among downregulated genes in neurons with *KIAA1429* knockdown compared to neurons expressing a non-targeting sgRNA. All depicted pathways have an adjusted P-value < 0.05.

Supplementary Fig 5. Neuronal cells with knockdown of genes essential for survival when forced to rely on glycolysis-only in high oxygen conditions (*MAPT, KIAA1429, TDRD3*) divert carbons from 13C-glucose to nucleotides (AMP, ADP, UDP), compared to neuronal cells expressing a non-targeting sgRNA. Knockdown of a less functionally related gene, *GLO1*, does not result in increased diversion of carbons derived from 13C-glucose to nucleotides, and actually decreases labeling in ADP and UDP metabolite pools. Data compiled from n = 3-4 samples per condition. *p < 0.05, **p < 0.01, ****p < 0.0001 by 2-way ANOVA with Šídák’s multiple comparisons test.

## Notes

### Competing Interest Statement

The authors have declared no competing interest.

